# Long-term offspring loss in lactating rats: Neurobiological and emotional consequences in a novel animal model

**DOI:** 10.1101/2025.02.17.634946

**Authors:** Luisa Demarchi, Alice Sanson, Anna-Lena Boos, Oliver J. Bosch

## Abstract

The maternal bond is a vital social connection that supports the survival and well-being of both the caregiver and offspring. Disruption of this bond, particularly following offspring loss, can result in profound trauma with long-lasting consequences. While considerable research has focused on the impact of maternal separation on offspring development, the biological effects of offspring loss on the mother remain largely unexplored. In this study, we examined the long-term effects of offspring loss on neuroplasticity, the oxytocin (OXT) and corticotropin-releasing factor (CRF) systems, and stress-coping behaviors in Sprague-Dawley rat mothers. We examined two groups of lactating mothers: (I) a control group, in which dams remained with their pups until natural weaning, and (II) a separated group, in which all offspring were removed on lactation day 1 and the mothers experienced offspring loss until the time corresponding to weaning (19 days).

Our results reveal that pup removal increased OXT receptor binding and reduced dendritic spine density in limbic brain regions, without altering estrogen receptor α or calbindin cell expression. Separated mothers additionally showed elevated plasma corticosterone levels and increased passive stress-coping behaviors in the forced swim test. Remarkably, passive stress-coping behavior was rescued by central CRF receptor blockade but not by OXT treatment, indicating that the CRF system plays a central role in the distress response to offspring loss. These findings establish the rat as a novel animal model for maternal distress, provide new insights into the complex neurobiology of grief, and suggests potential directions for future studies.

## Introduction

The maternal-infant bond is among the strongest social bonds in mammals^1^. In humans, disruption of this bond such as through offspring loss, elicits grief-like reactions in bereaved mothers, encompassing a range of emotional, cognitive, and behavioral responses, including depression and altered stress response. These dysregulations can progress to prolonged grief disorder (PGD)^2–4^. Despite the clinical relevance, the neurobiological consequences of offspring loss on the maternal brain -particularly in relation to neuroplasticity and the oxytocin (OXT) and corticotropin-releasing factor (CRF) systems-remain poorly understood. Rodent studies have been instrumental in characterizing maternal bonding. Reduced maternal care in rats negatively impacts the offspring development and acts as emotional trauma (for review see^5^). Importantly, repeated offspring separation has been shown to induce significant behavioral consequences in mothers^6-,9^. However, repeated separations do not fully model the experience of complete offspring loss -a scenario more analogous to human maternal grief. Only a limited number of studies have specifically investigated permanent offspring loss in rat mothers^10,11^ (for review see^12^). Our previous work demonstrates that offspring loss immediately after birth alters maternal brain function and stress-coping behavior during the first postpartum week, including heightened passive stress-coping, altered neuronal activity, and decreased OXT receptor (OXT-R) binding in the central amygdala (CeA)^13^. Building on these findings, we hypothesized that prolonged offspring loss over 19 days in rats could cause enduring neurobiological changes paralleling aspects of human grief^12^, with particular emphasis on the CRF and OXT systems. Disruption of social bonds in rodents affects neurotransmitter systems, neuroplasticity and behavior^14,15^. For example, in the monogamous and biparental prairie vole (*Microtus ochrogaster*) partner loss enhances CRF system signaling, reduces OXT signaling specifically in the nucleus accumbens (NAcc), and alters microglial activity and morphology in a brain region- and sex-specific manner^16–18^.

The CRF system, a key mediator of stress and depression, and has been extensively studied in both humans and rodents. Dysregulated CRF system activity is strongly linked to mood disorders^19–24^. CRF ligands (CRF, Urocortin 1-3) and their receptors (CRF-R1 and CRF-R2) are widely expressed both peripherally and centrally, and regulate physiological, autonomic, and behavioral responses to stress^20,25^. Elevated CRF levels in the cerebrospinal fluid and increased CRF-R binding have been reported in depressed patients^26^, consistent with upregulated CRF system activity^27,28^. In rats, CRF overexpression induces depressive-like symptoms, altered hypothalamic-pituitary-adrenal (HPA) axis function, and behavioral changes^29^, while chronic stress can lead to CRF system sensitization^30^. OXT, another key neuropeptide in social bonding, is essential for maternal bonding^31–33^, and other forms of social affiliation^34^. OXT-R are widely distributed throughout the brain, where OXT signaling modulates stress responses and facilitates social behavior^35^. OXT is thought to have antidepressant-like properties, partly by dampening HPA axis reactivity^36–38^, making it a candidate target for grief- and depression-related disorders^34,35^ including postpartum depression^39^.

Neuroplasticity is another important factor in grief and depression. Alterations in dendritic spine density, a key structural marker of plasticity, have been implicated in depression^40^, though the mechanisms remain incompletely understood^41^. Neuroplasticity and emotional processing are also regulated by estrogen receptor ESR1 by influencing dendritic spine formation and OXT-R expression^42–44^. ESR1 is expressed at high density in calbindin-immunoreactive (ir+) cells of the ventromedial hypothalamus (VMH)^45^, and calbindin-ir+ cells are reduced in the occipital cortex of post-mortem depressed patients^46^. We hypothesized that early loss of offspring would lead to enduring changes in the maternal stress-response, neuroendocrine systems and neuroplasticity. To test this, we studied lactating female Sprague-Dawley rats that experienced one day of motherhood followed by 19 days of offspring loss. We examined stress-coping behavior, HPA axis reactivity, OXT-R binding, dendritic spine density, and ESR1 and calbindin ir+ cells expression in key limbic and maternal brain regions. This work provides novel insights into the neurobiology of maternal distress and highlights the potential therapeutic relevance of targeting central CRF-R signaling.

## Materials and Methods

### Animals

Sprague-Dawley rats were obtained from Charles River Laboratories (Sulzfeld, Germany) and kept under standard laboratory conditions (12:12h light/dark cycle, lights on at 7 a.m., room temperature 22 ± 1°C, relative humidity 55 ± 5°C) with access to standard rat chow (ssniff-Spezialdiäten GmbH, Soest, Germany) and water *ad libitum*. After 7 days of habituation, females were mated with sexually experienced male Sprague-Dawley rats in standard laboratory cages (Eurostandard Type IV, 60 cm x 40 cm x 20 cm) for 10 days. From pregnancy day 18 on, pregnant females were single-housed for undisturbed delivery in observational cages (plexiglass, 38 cm x 22 cm x 35 cm); all rats delivered 8-15 pups within 3-4 days after being single-housed. The studies were conducted in accordance with the ARRIVE guidelines, the European regulations of animal experimentation (European Directive 2010/63/EU) and were approved by the local government of Unterfranken (Bavaria, Germany). According to the 3-Rs principles, all efforts were made to minimize the number of animals and their distress or suffering.

### Study design

We investigated two primary groups of lactating females: (I) control mothers, which remained with their pups until natural weaning, and (II) separated mothers, which had all offspring removed on lactation day 1 and experienced offspring loss for 19 days (**Fig. 1**). The 19-day separation period was selected because it corresponds to the pre-weaning stage, a critical time when the mother–offspring bond remains strong in rats^47^. Our study followed a stepwise approach. First, we assessed the effects of prolonged offspring loss on OXT-R binding, ESR1 and calbindin-immunoreactive (ir⁺) cells in brain regions implicated in maternal behavior. This analysis provided an initial characterization of neuroendocrine adaptations within the maternal brain and helped identify target regions for subsequent investigation. Next, we evaluated dendritic spine density and behavioral outcomes, incorporating an additional group of *virgin* females to distinguish experience-dependent plasticity in maternal neural circuits.

**Figure 1:**
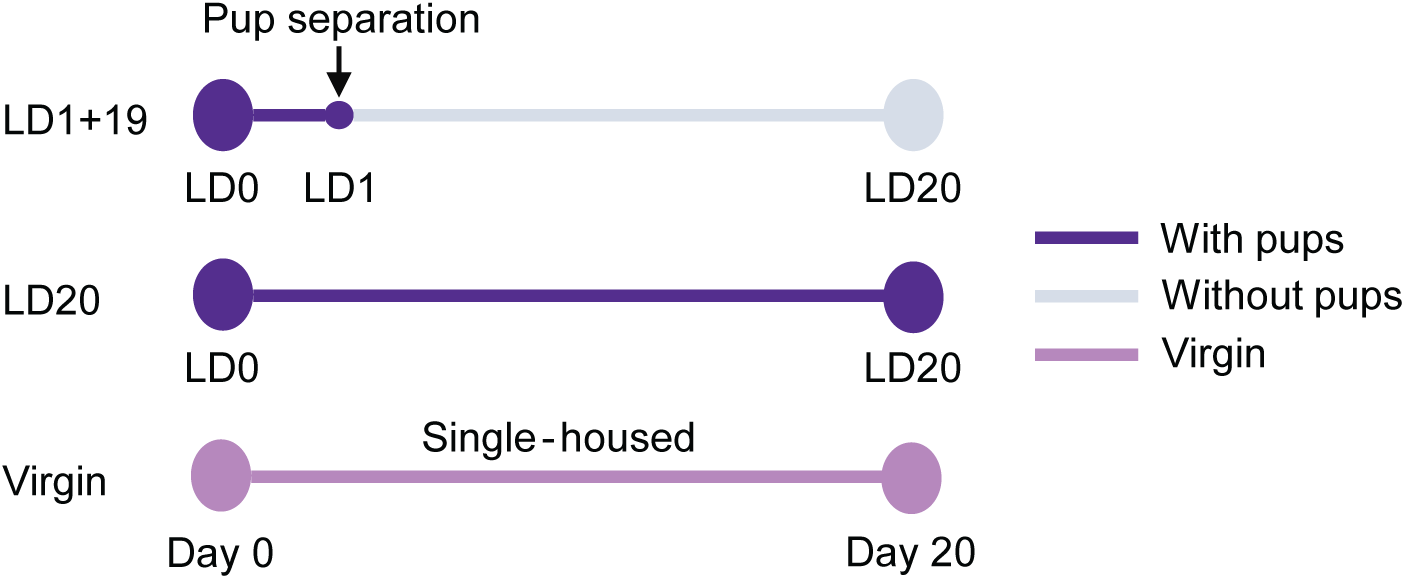
Schematic timelines summarizing the maternal separation paradigm. Abbreviations: LD, lactation day.

### Experimental groups

Rats were randomly assigned to one of three experimental groups: (I) control lactating mothers (primiparous) that remained with their pups until weaning (lactation day 20, *LD20* group), (II) separated mothers (*LD1+19* group; primiparous) whose litters were removed on lactation day 1 (LD1), and (III) *virgin* females (*virgin* group; nulliparous). To standardize litter size, each litter was adjusted to eight pups of mixed sexes on the day of delivery (LD0). In the *LD1+19* group, all pups were removed on LD1 at 10:00 a.m., after which the mothers were left undisturbed in their home cages. To control for the effects of single housing, *virgin* females were housed individually for the same duration as separated mothers.

### Experimental design

*Experiment 1*: a first cohort of rats followed the protocol described above and was sacrificed on lactation day 20 (LD20) without further manipulations (**Fig. 1**). Brains were collected for: (i) OXT-R binding analysis (*LD20* = 6; *LD1+19* = 6), (ii) immunofluorescence analysis (*LD20* = 7; *LD1+19* = 5), and (iii) dendritic spine analysis (*LD20* = 4; *LD1+19* = 4; *virgin* = 5).

*Experiment 2:* a second cohort (*LD20* = 15; *LD1+19* = 15; *virgin* = 9) was tested for locomotor and anxiety-like behavior in the light–dark box (LDB) on LD19 and in the open field (OF) on LD20. Immediately following the final behavioral test, rats were sacrificed. *Experiment 3*: a third cohort (*LD20* = 13; *LD1+19* = 14; *virgin* = 8) was evaluated for stress-coping behavior in the single-session forced swim test (FST) on LD20. Within 5 minutes of test completion, rats were sacrificed, and trunk blood was collected for plasma corticosterone (CORT) analysis.

*Experiments 4 and 5*: two additional cohorts were prepared following the same protocol. On LD15, all rats underwent stereotaxic implantation of an intracerebroventricular (icv) guide cannula for acute substance infusion. On LD19, they completed the 15-min pre-test session of the two-session forced swim test (FST). On LD20, they were tested in a 10-min session following infusion of either vehicle (VEH) or a pharmacological treatment. In Experiment 4, the CRF-R1/2 antagonist D-Phe was administered (*LD20 VEH* = 10; *LD1+19 VEH* = 8; *LD1+19 D-Phe* = 9), whereas in Experiment 5, synthetic OXT was infused (*LD20 VEH* = 8; *LD1+19 VEH* = 8; *LD1+19 OXT* = 7). In both experiments, all rats were sacrificed immediately after the final behavioral test and brains taken to verify cannula placement.

### Brain analyses

#### OXT-R autoradiography

Rats were anesthetized and underwent cardiac perfusion with ice-cold 1x PBS and were decapitated. Brains were flash-frozen in n-methylbutane, and stored at -20 °C until cutting into coronal sections of 16 μm using a cryostat (CM3050S, Leica Microsystem GmbH). For each brain region of interest (bed nucleus of the stria terminalis (BNST), nucleus accumbens shell (NAcc shell), medial preoptic area (MPOA), accessory olfactory bulbs (AOB), central amygdala (CeA), agranular insular cortex (AIP), ventromedial hypothalamus (VMH), lateral septum (LS), prelimbic cortex (PL)) six slices per rat were collected on SUPERFROST microscope slides and stored at -20 °C until further processing. The OXT-R autoradiography was performed following an established protocol^48^. Briefly, the ornithin-vasotocin analogue [125I]-OVTA [d(CH2)5[Tyr(Me)2,Thr4,Orn8,[125I]Tyr9-NH2]; Perkin Elmer, USA) was used as a tracer. First, the slides were thawed and allowed to dry thoroughly at room temperature. The tissue was shortly fixed via 0.1 % PFA, washed 2 x in Tris (50 mM, pH 7.4), covered with the tracer solution (50 mM tracer, 10 mM MgCl2, 0.1 % BSA) for 60 min, washed 3 x in Tris / MgCl2 buffer for 7 min, each, followed by 30 min spinning in Tris / MgCl2. Finally, slides were dipped into water and air dried before being exposed to Biomax MR films (Kodak, Cedex, France) for 15 days. The films were scanned using the EPSON Perfection V800 Scanner (Epson GmbH, Munich, Germany), and the optical density of each region of interest was analyzed using ImageJ^49^ by subtracting the background activity as previously described^48,50^. The analyses were performed simultaneously for 6 slices per rat and per region.

#### Immunofluorescence staining

Rats were anesthetized and perfused with ice-cold 1x PBS and subsequently with 4 % paraformaldehyde (PFA) dissolved in 1x PBS. Brains were removed, fixed overnight in 4 % PFA, and subsequently incubated in 30 % sucrose in 1x PBS until the brain sank. After fixation and cryoprotection in sucrose, brains were flash-frozen in n-methylbutane, and stored at -20 °C until cutting into coronal sections of 16 μm using a cryostat (CM3050S, Leica Microsystem GmbH, Nussloch, Germany). Six consecutive slices containing the VMH region were collected per rat on SUPERFROST microscope slides and stored at -20 °C until further processing. Slides were washed in 1x PBS, permeabilized with 0.3 % Triton, blocked with 5 % normal goat serum (Vector Laboratories, Newark, USA), and incubated overnight at 4 °C with primary antibodies (Merck Millipore, rabbit anti ER alpha 1:2000; Merck, mouse anti-NeuN 1:1000; Synaptic Systems, chicken anti-calbindin 1:2000). The next day, slides went through 1 h incubation with secondary antibodies (Alexa Fluor 488-conjugated anti-rabbit IgG 1:800, Alexa Fluor 594-conjugated anti-mouse IgG 1:800 and Alexa Fluor 649-conjugated anti-chicken IgG 1:800) at room temperature. All slides were finally mounted using ROTI®Mount FluorCare DAPI (Carl Roth, Karlsruhe, Germany).

#### Golgi staining for spine density visualization

The spine density was assessed in *virgin*, *LD20* and *LD1+19* groups to investiganeuronal plasticity. Firstly, rats were sacrificed with CO_2_ and brains were collected. The Rapid GolgiStainTM Kit (FD NeuroTechnologies, Columbia, USA) was used and applied according to the manufacturer’s protocol. Brains were cut into 100 μm slices using a cryostat (CM3050S, Leica Microsystem GmbH) at -28° and placed on gelatin-coated microscope slides (FD NeuroTechnologies, Columbia, USA). After complete drying, Golgi staining was performed, and sections were stored at room temperature in the dark before acquiring images using a brightfield microscope (Leica Thunder DM6 B, Camera Leica DFC9000 GT). ImageJ^49^ was used to count the spines of secondary dendrites in the PL, AIP, BLA and VMH. Secondary dendrites from different neurons (between 4 and 12, depending on the brain region) were chosen from 10X overview preview images followed by acquiring 40X focus images for analysis. The ROI of each dendrite was selected, brightness and sharpening function were adjusted. Spines were counted on a 25-30 μm^2^ section of the dendrite (each dendrite belonging to a different neuron) and the absolute spine density was calculated by using the following formula^51^: (n = number of spines; μm = dendritic length): 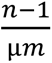.

### Behavioral experiments

#### Light dark box (LDB)

On LD19, between 9:00 a.m. and 12:00 p.m., rats from experiment 2 were tested in the LDB to assess anxiety-like behavior^52^. The apparatus consisted of a light compartment (40 × 50 cm, 180 lux) and a dark compartment (40 × 30 cm, 0 lux) connected by a 7.5 × 7.5 cm opening. At the start of the test, rats were placed in the center of the light box, and behavior was recorded for 10 min using EthoVision XT (Noldus, Wageningen, the Netherlands) by an experimenter blind to group assignment. The following parameters were analyzed: (a) percentage of time spent in the light box, (b) latency to re-enter the light compartment, and (c) locomotor activity. The arena was cleaned with tap water and dried thoroughly between trials.

#### Open-field (OF)

On LD20, between 9:00 a.m. and 12:00 p.m., rats from experiment 2 were tested in the OF to evaluate locomotor activity and anxiety-like behavior^53^. The apparatus was an empty rectangular arena (80 × 80 cm). Rats were placed in one corner and allowed to explore freely for 10 min. Behavior was recorded using EthoVision XT, with analysis performed by an experimenter blind to treatment. The following measures were scored: (a) locomotion, (b) velocity, and (c) number of center entries. The arena was cleaned with tap water and dried between trials.

#### Forced swim test (FST)

Two versions of the FST were employed to assess passive stress-coping behavior. *Rationale:* we used a single-session protocol to capture baseline group differences without pre-exposure, whereas the two-session protocol provided higher sensitivity to drug effects -particularly relevant because animals in experiment 4-5 underwent surgery (see below).

#### Single-session FST

On LD20, between 9:00 a.m. and 12:00 p.m., rats from experiment 3 were tested in a single 10-min FST session. Each rat was placed in a cylindrical tank (50 cm high, 30 cm diameter) filled with tap water (23 ± 1 °C) at a depth preventing contact with the bottom. Behavior was recorded and analyzed using JWatcher (https://www.jwatcher.ucla.edu) by an experimenter blind to group assignment. The total time spent immobile (floating) was quantified as an index of passive stress-coping, which is reminiscent of depressive-like behavior^54^.

#### Two-session FST

In experiments 4 and 5, rats were tested using the two-day Porsolt FST, the standard paradigm for evaluating antidepressant treatments^54^. These animals had previously undergone stereotaxic surgery (see details below) and were tested 5 days postoperatively. On LD19, rats underwent a 15-min pre-test session. On LD20, they were exposed to a 10-min test session following intracerebroventricular infusion of vehicle or a pharmacological agent. The two-session design was chosen to stabilize baseline immobility at the pre-test and to maximize the dynamic range for detecting drug-induced reductions at the test while controlling for post-surgical variability. All other experimental conditions and behavioral analyses were identical to those described for the single-session FST.

### Stereotaxic surgery

On LD15, rats of experiments 4 and 5 were implanted with a stainless-steel guide cannula (21-G, length 12 mm), above the right ventricle (1.0 mm posterior, 1.6 mm lateral to bregma, 1.8 mm ventral)^55^. Surgeries were performed under isoflurane anesthesia, following previously described procedures^56^. The guide cannula was closed using a 25-G stainless steel stylet to prevent occlusion. To minimize post-surgical discomfort, all rats received a subcutaneous injection of buprenorphine (0.05 mg/kg, Bayer Vital GmbH, Leverkusen, Germany) prior to surgery. Following surgery, rats were monitored continuously until recovery from anesthesia and then returned to their home cages. Rats were observed daily for general health, body weight, and signs of discomfort or infection throughout the recovery period.

### Drug administration

On LD20, rats of experiment 4 and 5 received a single acute icv infusion 10 min prior to the second session of the two-day forced swim test. Depending on the treatment group, animals were infused with vehicle (VEH; 5 µL sterile Ringer’s solution, pH adjusted to 7.4; Braun, Melsungen, Germany), the human/rat CRF-R1/2 antagonist D-Phe [(D-Phe¹², Nle²¹–³⁸, α-Me-Leu³⁷)-CRF (12–41); 10 µg/5 µL; Bachem, Bubendorf, Switzerland], or synthetic OXT (1 µg/5 µL; Tocris, Nordenstadt, Germany). All doses were selected based on previous studies^56^. Infusions were performed using a 25 G stainless-steel infusion cannula (14 mm length) connected to a 10 µL Hamilton micro syringe via PE-50 tubing (50 cm). The infusion cannula was inserted into the pre-implanted guide cannula and held in place with a piece of silicon tubing. Substances were delivered over 30s, after which the cannula was left in position for an additional 30s to minimize reflux before removal, as previously described^38^.

### Verification of cannula placement

Rats of experiment 4 and 5 were sacrificed immediately following the last behavioral test with CO_2_ and blue dye was injected via the infusion system into the guide cannula. Verification of the infusion site of the icv cannula was observed by dye spread throughout the ventricular system. Only animals with correct verified infusion sites were included in the statistical analyses.

### ELISA for plasma CORT

In experiment 3, rats were sacrificed within 5 min after the single-session FST, and approximately 1 mL of trunk blood was collected in EDTA-coated tubes on ice (Sarstedt, Numbrecht, Germany). Samples were centrifuged at 4 °C (10,000 rpm/12298 rcf) for 10 min. Plasma was aliquoted and stored at −20 °C until analysis. CORT concentrations were quantified using a commercially available ELISA kit (Tecan IBL International GmbH, Hamburg, Germany) following the manufacturer’s instructions. All samples were measured in duplicate, and mean values were used for statistical analysis.

### Statistical analysis

All statistical analyses were performed with GraphPad PRISM 9 (GraphPad Software, San Diego, USA). Data were first tested for normality (Shapiro-Wilk or Kolmogorov– Smirnov test) and homogeneity of variance (Brown-Forsythe test). Outliers were identified using the ROUT method.

Group comparisons of OXT-R binding, ESR1 and calbindin-ir+ cell counts were performed using two-sample Student’s *t*-test. Dendritic spine density, plasma CORT concentration, anxiety-like behavior, locomotor activity and stress-coping behavior were analyzed by one-way ANOVA followed by Sidak’s post hoc multiple comparisons. Data are expressed as mean ± SEM; and as significance was set at *p* ≤ 0.05.

## Results

### Increased OXT-R binding in the VMH and PL after offspring loss

To determine whether offspring loss alters OXT-R binding within maternal and limbic brain regions, we performed receptor autoradiography in *LD20* and *LD1+19* mothers. *LD1+19* dams exhibited significantly higher OXT-R binding in the ventromedial hypothalamus (VMH; *M* = 0.257, *SD* = 0.077) compared to *LD20* dams (*M* = 0.147, *SD* = 0.083; *t*(10) = 2.366, *p* = 0.039; **Fig. 2A**). A similar finding was observed in the prelimbic cortex (PL), where OXT-R binding was slightly elevated in *LD1+19* (*M* = 0.070, *SD* = 0.005) relative to *LD20* dams (*M* = 0.066, *SD* = 0.002, *t*(10) = 2.203, *p* = 0.052; **Fig. 2B**).

**Figure 2:**
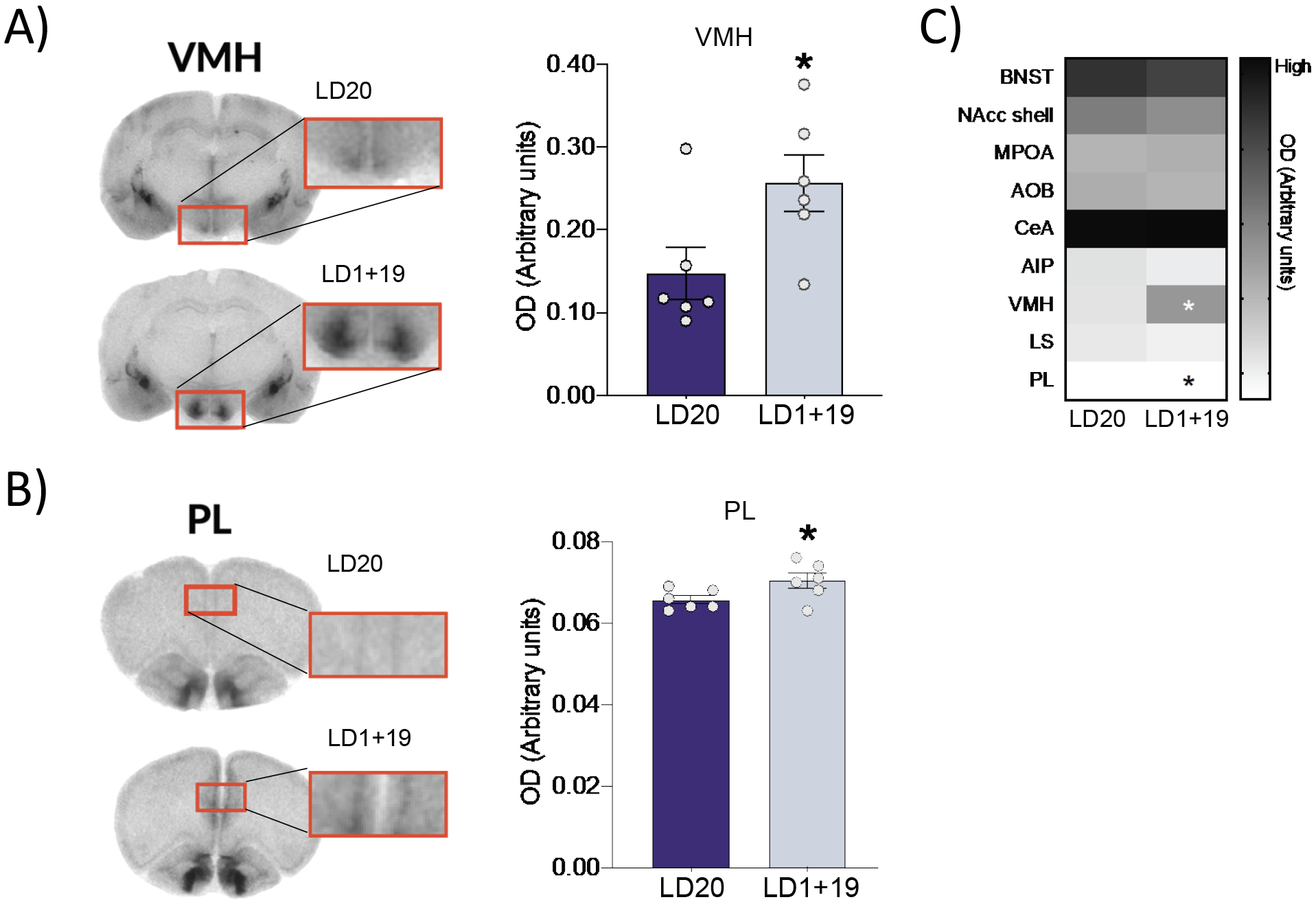
OXT-R binding in brain regions of the limbic system and maternal network. (**A,B**) Representative coronal brain sections of the VMH and PL showing differences in OXT-R binding. Optical density (arbitrary units) for OXT-R binding in the (**A**) VMH and (**B**) PL. (**C**) Overview heatmap of gray density (arbitrary units) for OXT-R binding in the analyzed brain areas (* p ≤ .05 versus *LD20*). Abbreviations: AIP, agranular insular cortex; AOB, accessory olfactory nuclei; BNST, bed nucleus of the stria terminalis; CeA, central amygdala; LS, lateral septum; MPOA, medial preoptic area; NAcc, nucleus accumbens; PL, prelimbic cortex; VMH, ventral medial hypothalamus. Student’s t Test. Data are expressed as mean ±SEM. * p ≤ .05 versus *LD20*.

No significant group differences were detected in the other regions analyzed (**Fig. 2C**): MPOA (*t*(10) = 0.215, *p* = 0 .834); BNST (*t*(10) = 0.828, *p* = .427); CeA (*t*(10) = 0.061, *p* = 0.952); OB (*t*(10) = 0.595, *p* < 0 .565); NAcc shell (*t*(10) = 1.440, *p* = 0 .180); AIP (*t*(10) = 1.212, *p* = 0 .254); LS (*t*(10) = 0.613, *p* = 0 .554)].

### ESR1 and calbindin ir+ cells in the VMH and PL did not differ between groups

To investigate whether the increased OXT-R binding in the VMH and PL (see above) was associated with changes in ESR1 expression, we quantified ESR1- and calbindin-immunoreactive (ir+) cells in the VMH and PL of *LD20* and *LD1+19* mothers. In the VMH, no significant group differences were detected in the proportion of ESR1-ir+ cells (*t*(10) = 0.456, *p* = 0.658; **Fig. 3A, B**) or of calbindin ir+ cells (*t*(10) = 0.706, *p* = 0.496; **Fig. 3A, C**). ESR1-ir+ cells were not detectable in the PL (data not shown).

**Figure 3:**
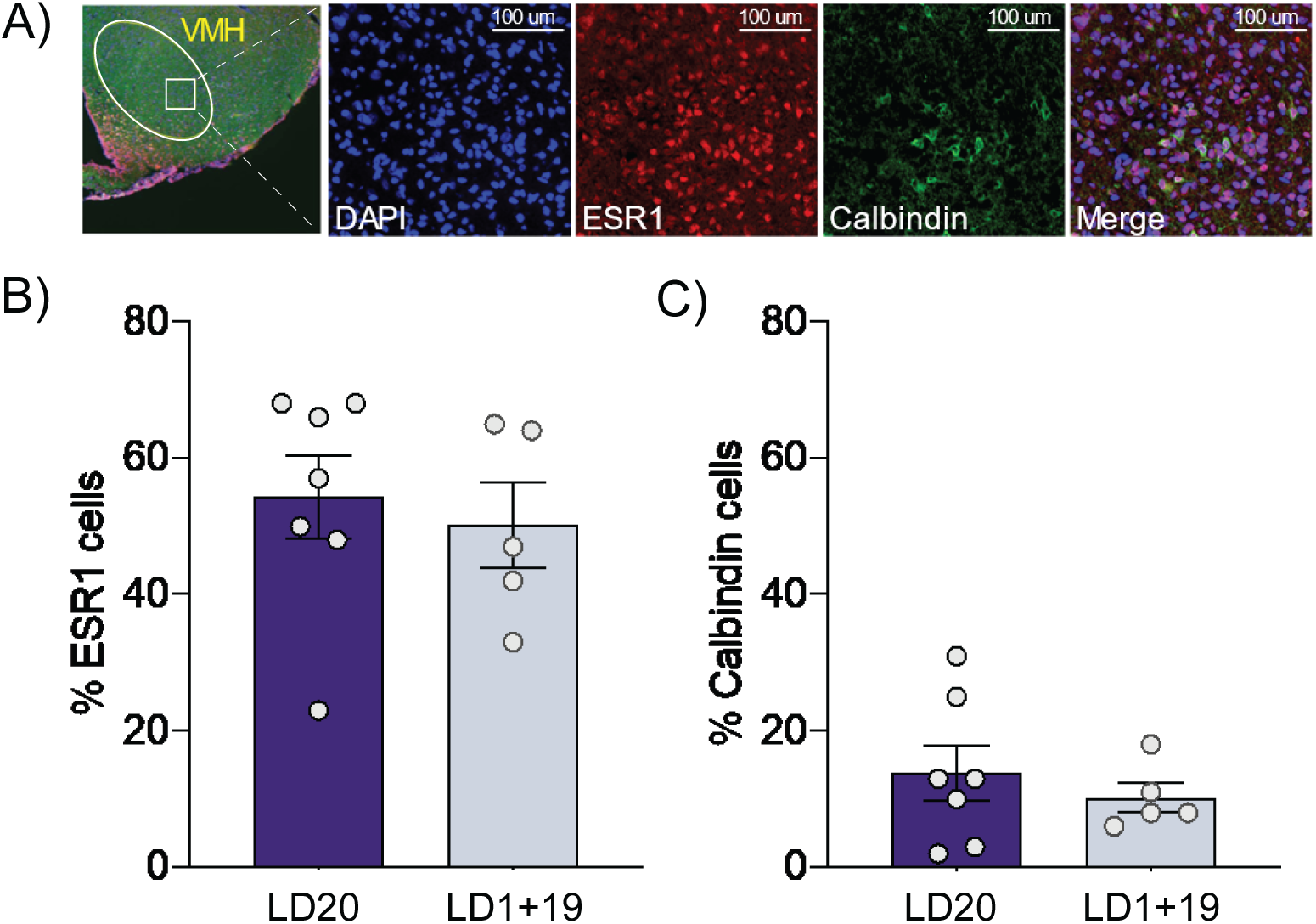
ESR1 and calbindin ir+ cells in the VMH. (**A**) Representative fluorescent images showing DAPI (blue), ESR1 cells (red), calbindin (green) and merged channels in the VMH. Percentage of (**B**) ESR1 and (**C**) calbindin ir+ cells in the VMH. One-way ANOVA. Data are expressed as mean ± SEM.

### Reduced secondary dendritic spine density in the VMH after offspring loss

To assess the impact of offspring loss on neuroplasticity, we quantified secondary dendritic spine density in the VMH, PL, BLA and AIP of *virgin*, *LD20* and *LD1+19* females (**Fig. 4A**), based on regions identified in our previous study^13^. Statistical analysis revealed a significant group effect in the VMH (one-way ANOVA: *F*(2, 10) = 8.734; *p* = 0.006; **Fig. 4B**). Post hoc analyses showed that *LD1+19* dams exhibited significantly lower spine density compared to *virgin* (*p* = 0.024) and *LD20* mothers (*p* = 0.009). In contrast, no significant differences were observed in the other regions examined. No differences in spine density were found for the PL (*virgin*: 0.64 ± 0.07; *LD20*: 0.53 ± 0.04; *LD1+19* : 0.62 ± 0.04; one-way ANOVA: *F*(2,15) = 0.590, *p* = 0.356), the BLA (*virgin*: 0.61 ± 0.03; *LD20*: 0.64 ± 0.009; *LD1+19*: 0.61 ± 0.02; one-way ANOVA: *F*(2,16) = 0.158, *p* = 0.855), and the AIP (*virgin*: 0.62 ± 0.05; *LD20*: 0.69 ± 0.04; *LD1+19*: 0.68 ± 0.04; one-way ANOVA: *F*(2,15) = 0.599, *p* = 0.562).

**Figure 4:**
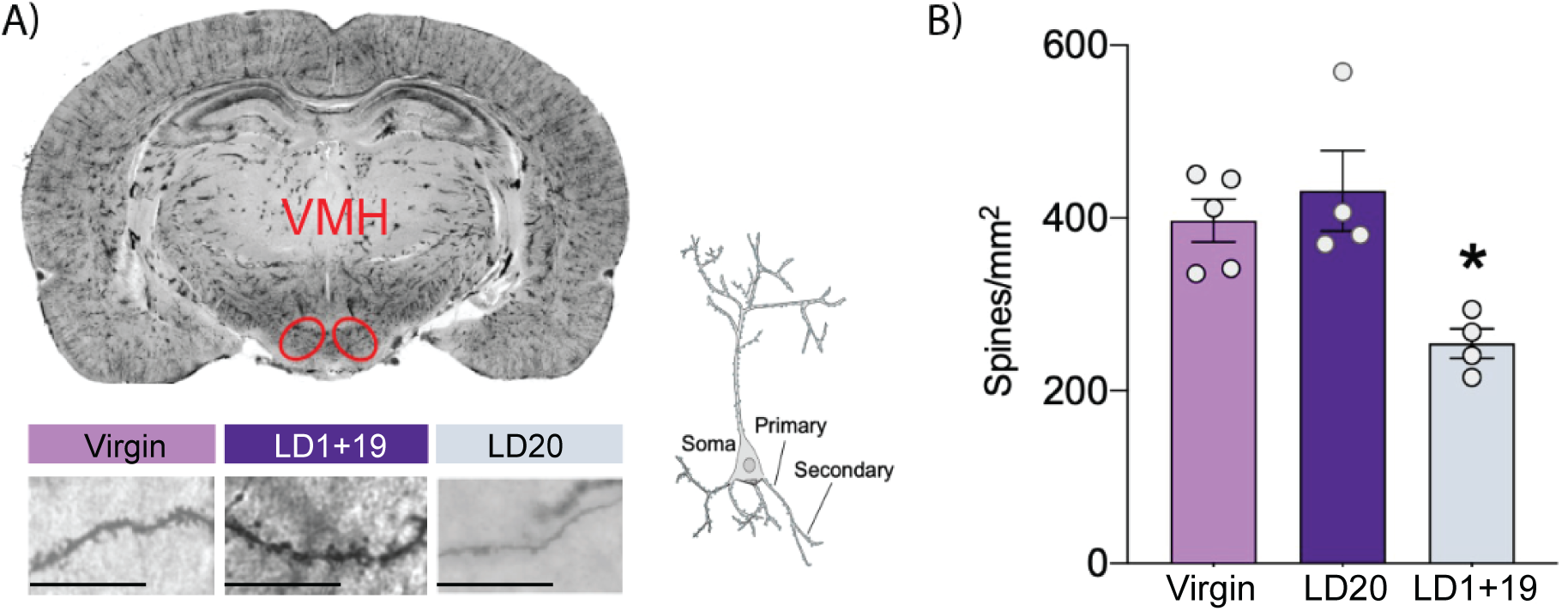
Secondary dendritic spine density in the VMH. (**A**) Representative coronal whole brain section with Golgi staining and schematic illustration of the neuronal dendritic arbor (Scale bar 10 µm). (**B**) Absolute spine density / mm^2^ across experimental groups. (**C**) One-way ANOVA. Data are expressed as mean ± SEM. * p < .05 versus all other groups.

### Anxiety-like behavior was not affected by offspring loss

To examine whether offspring loss altered anxiety-like behavior, *virgin*, *LD20*, and *LD1+19* rats were tested in the LDB on LD19. Groups differed in latency to re-enter the light compartment (Kruskal–Wallis test, *p* = 0.045), with *LD20* dams tending to be faster (48.6 s ± 15.9 s) than *virgin* females (213.5 s ± 77.5 s; Dunn’s test, *p* = 0.058). *LD1+19* dams did not differ from either group (121.9 s ± 31.9 s).

No group differences were observed in the time spent in the light compartment (*virgin*: 34.0 s ± 6.3 s; *LD20*: 36.6 s ± 3.6 s; *LD1+19*: 42.0 s ± 5.1 s; one-way ANOVA: *F*(2,36) = 0.676, *p* = 0.515) or in locomotor activity within the LDB (*virgin*: 3778 cm ± 326 cm; *LD20*: 3733 cm ± 194 cm; *LD1+19*: 3851 cm ± 312 cm; one-way ANOVA: *F*(2,36) = 0.054, *p* = 0.948). Thus, *LD1+19* mothers did not differ from either *virgin* or *LD20* mothers in any measure of anxiety-like behavior.

### Locomotor activity was not affected by offspring loss

The same cohort of rats was tested in the OF on LD20. Groups differed significantly in distance travelled (one-way ANOVA: *F*(2,36) = 3.773, *p* = 0 .032; **Fig. 5A**) and in velocity (one-way ANOVA; [*F*(2,36) = 3.788], *p* = 0.032; **Fig. 5B**). Post hoc comparisons revealed that *virgin* females showed greater locomotion (*p* = 0.029) and velocity (*p* = 0.028) compared with *LD20* dams. *LD1+19* mothers did not differ from either group. No significant differences were found in the number of center entries (*virgin*: 21.4 ± 2.7; *LD20*: 16.3 ± 1.5; *LD1+19*: 20.5 ± 2.0; one-way ANOVA: *F*(2,36) = 1.858, *p* = 0.170). Overall, offspring loss did not alter locomotor activity.

**Figure 5:**
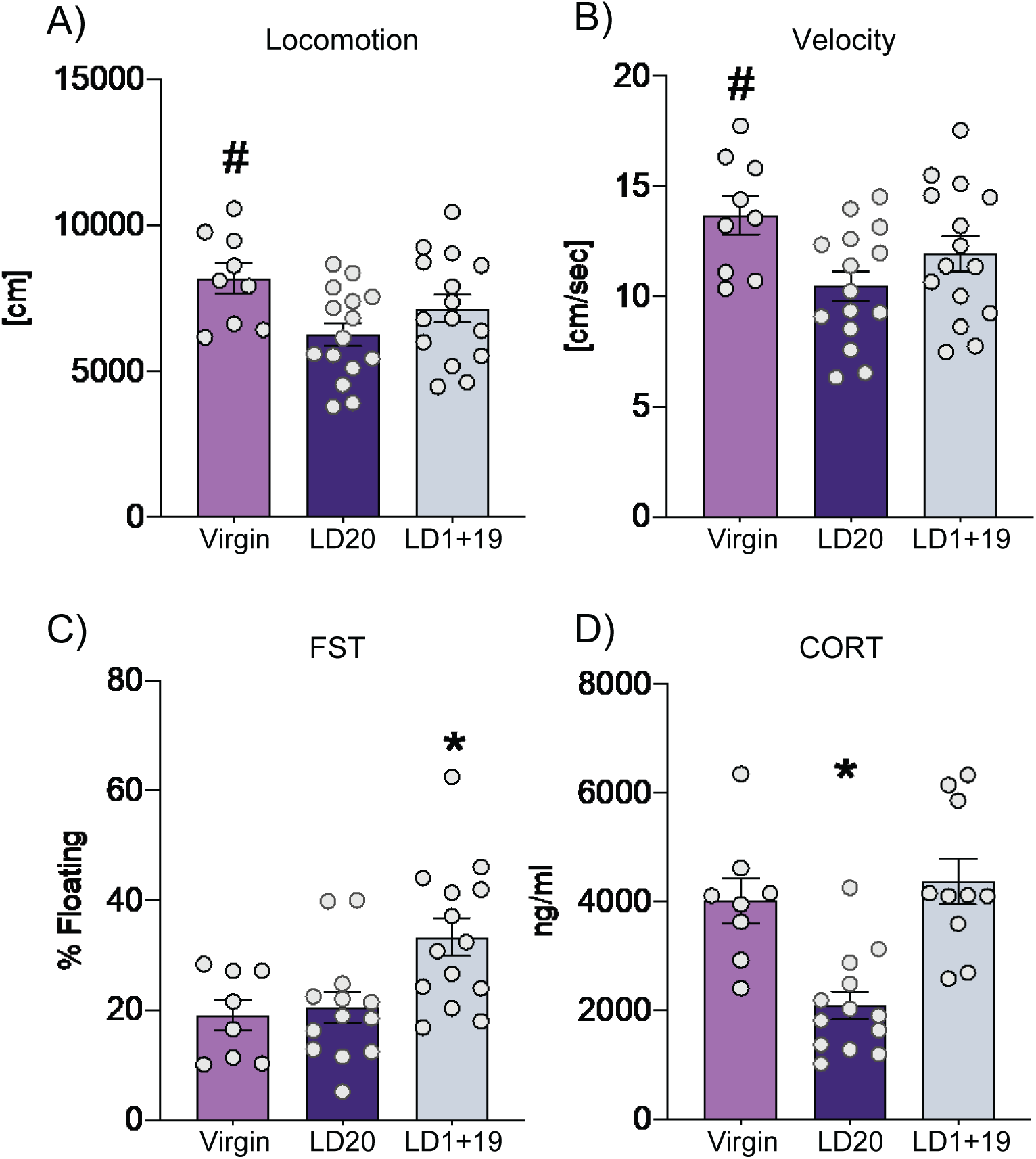
(**A)** Locomotor behavior and **(B**) velocity in the OF, (**C**) passive stress-coping behavior in the classic FST, and (**D**) corresponding plasma CORT concentration. (**A**) Total distance traveled in cm and (**B**) mean velocity over the 10-min OF test. (**C**) Percentage of floating during the 10-min classic FST. (**D**) Plasma CORT concentration 15 min after start of the FST. One-way ANOVA. Data are expressed as mean ± SEM. # p < .05 versus *LD20 VEH*; * p < .05 versus all other groups.

### Offspring loss increased passive stress-coping behavior and induced a virgin-like stress response

A different cohort of *virgin*, *LD20* and *LD1+19* rats was tested for passive stress-coping behavior in the single-session FST on LD20. Groups differed significantly in time spent floating (one-way A N O VA : *F*(2,32) = 6.233, *p* = 0.005; **Fig. 5C**). *LD1+19* dams spent more time floating compared with both *virgin* (*p* = 0.019) and *LD20* (*p* = 0.014) rats. Stress-induced plasma CORT levels, which were collected 15 min after the start of the FST, also differed across groups (*F*(2,28) = 13.50, *p* = 0.0001; **Fig. 5D**). Post hoc comparisons revealed significantly lower CORT concentrations in *LD20* compared with *virgin* (*p* = 0.002) and *LD1+19* (*p* = 0.0001) rats. *LD1+19* mothers did not differ from *virgin* females (*p* > 0.05), suggesting that offspring loss abolished the lactation-induced blunting of HPA axis activity.

### Central CRF-R1/2 blockade normalized passive-stress coping behavior in *LD1+19* mothers

Following a pre-test on LD19, *LD20* and *LD1+19* rats were tested in the FST on LD20 after ICV infusion of VEH or the CRF-R1/2 antagonist D-Phe. Groups differed significantly in floating behavior (one-way ANOVA: *F*(2,23) = 5.828, *p* = 0.013; **Fig. 6A**). *LD1+19 VEH* mothers spent more time floating than *LD20 VEH* rats (*p* = 0.047), replicating the findings from the single-session FST (**Fig. 5C**). Importantly, floating time was significantly lower in *LD1+19 D-Phe* rats compared to *LD1+19 VEH* (*p* = 0.014), reaching levels indistinguishable from *LD20 VEH* (*p* > 0.05).

**Figure 6:**
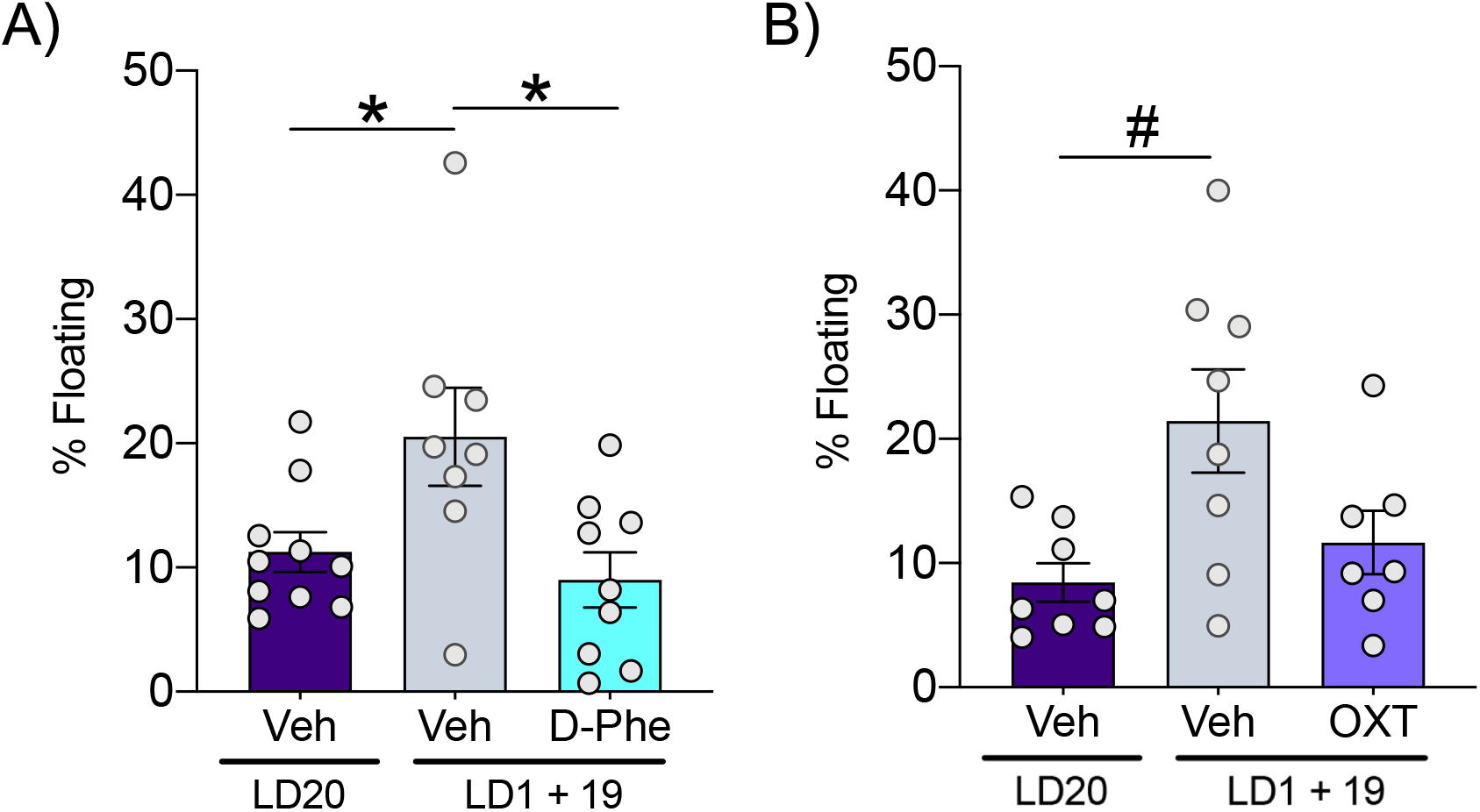
(**A**) Passive stress-coping behavior in the FST following central pharmacological manipulation of the brain CRF and (**B**) OXT systems. (**A**) Percentage of floating during the 10-min modified FST following icv infusion of VEH or the CRF-R1/2 antagonist D-Phe, and (**B**) of VEH or synthetic OXT. One-way ANOVA. Data are expressed as mean ± SEM. * p < .05 versus all other groups; # p < .05 versus *LD20 VEH*.

### Central OXT infusion did not alter passive-stress coping behavior in *LD1+19* mothers

In a different cohort, *LD20* and *LD1+19* rats were tested in the FST after infusion of VEH or synthetic OXT. Groups differed in floating behavior (one-way ANOVA: *F*(2,20) = 5.273, *p* = 0.014; **Fig. 6B**). Consistent with previous experiments, *LD1+19 VEH* rats floated more than *LD20 VEH* (*p* = 0.014). However, infusion of OXT in *LD1+19* mothers was not sufficient to alter floating when compared with *LD1+19 VEH* (*p* = 0.08).

## Discussion

The loss of offspring represents a profound life event with strong effects on physiology and behavior. In humans, such experiences are associated with marked changes in emotional and physical health, including a higher risk for prolonged grief disorders^4,57–59^. To investigate the underlying mechanisms and to identify potential therapeutic targets, appropriate animal models are required. In this study, we build upon our previous work on consequences of short-term separation on rat mothers^13^. We examined the long-term consequences of offspring loss and demonstrate that extended separation manifests in distinct neurobiological and behavioral adaptations. After 19 days of pup removal, mothers exhibited increased OXT-R binding specifically in the VMH and PL, had reduced dendritic spine density in the VMH, and heightened HPA axis reactivity together with increased passive stress-coping behavior. Importantly, central pharmacological blockade of CRF-R1/2 normalized the impaired behavior, whereas OXT infusion did not, highlighting the prominent role of CRF system activity in mediating maternal distress responses due to pup loss.

### Increased OXT-R binding in the VMH and PL after offspring loss

During the peripartum period, the maternal brain undergoes dynamic adaptations in OXT signaling that support the initiation and maintenance of maternal behaviors^60,61^. In our study, OXT-R binding was significantly elevated in the VMH and PL following offspring loss, contrary to our initial hypothesis that separation would reduce OXT signaling. The VMH, which contains a high density of OXT-Rs and plays a role in maternal care, aggression, and defensive behaviors^62–66^, has not been studied in the context of pup removal. The PL, a region critical for cognitive and emotional regulation^67,68^, has also been implicated in maternal care, as OXT-R blockade in this area disrupts caregiving behavior^69^. Thus, the upregulation of OXT-Rs in both the VMH and PL may reflect adaptive—yet potentially insufficient—neurobiological adjustments triggered by permanent offspring removal. Previous studies investigating repeated maternal separation also reported changes in OXT-R binding in the PL and MPOA^6,70,71^, consistent with the sensitivity of OXT signaling to maternal experience. Reduced maternal interactions, including fewer opportunities for the milk ejection reflex, likely decrease endogenous OXT release^72^, which has been associated with reduced activity of OXT-positive neurons in the PVN^73^. Based on this evidence, we propose that permanent offspring loss alters OXT dynamics, with potentially less OXT release leading to the detected OXT-R upregulation thereby serving as a compensatory adaptation. Future studies should examine brain region-specific OXT release and transcriptional regulation to provide a more complete picture of system-wide neuroendocrine adaptations.

### ESR1 and calbindin do not mediate OXT-R changes

Estrogen has been shown to regulate OXT-R expression via ESR1 and to interact with calbindin^44,74–76^. To assess whether the OXT-R upregulation observed in our study could be explained by alterations of these factors, we quantified ESR1- and calbindin-immunoreactive (ir⁺) cells in the VMH and PL. No differences were detected in the proportion of ESR1- or calbindin-ir⁺ cells in the VMH between groups, and ESR1 was not detectable in the PL. These results indicate that the increase in OXT-R binding following offspring loss is unlikely to be mediated by changes in ESR1 or calbindin expression. Instead, other neuroendocrine pathways may contribute to the observed receptor alterations. In particular, CRF signaling represents a likely candidate, given its known interactions with the OXT system and its role in stress-related neuroadaptations^77^.

### Reduced spine density in the VMH after offspring loss

Neuroplasticity is a central component of maternal adaptations, with dendritic spine remodeling occurring across pregnancy and the postpartum period^78^. In our study, we observed a selective reduction in secondary dendritic spine density within the VMH after offspring loss, whereas no significant changes were detected in other regions examined. Alterations in dendritic spine density have been widely reported in response to stress and depression in both humans and rodents^79–89^. Given the VMH’s established role in the regulation of stress reactivity, social interactions, and aggression^90–92^, the observed decrease in spine density may contribute to the stress-response dysregulation observed in separated mothers. Importantly, these structural changes were accompanied by increased OXT-R binding in the VMH, indicating that both cellular and receptor-level adaptations converge in this region. A plausible interpretation is that reduced spine density may reflect altered excitatory connectivity within VMH circuits, while the concurrent increase in OXT-R binding could shift the balance of neuromodulatory input, thereby reshaping local network responsiveness. Rather than acting as a simple compensatory mechanism, these parallel adaptations may represent a reorganization of VMH function following offspring loss, potentially influencing how this region integrates social and stress-related signals. A limitation of the present work is that estrous cycle phases of *virgin* females were not monitored, which could have influenced spine density measurements. Future studies should therefore control for reproductive state and directly test the functional link between OXT-R signaling and structural remodeling in the VMH to clarify how these adaptations interact to shape maternal behavioral responses.

### Anxiety-like and locomotor behaviors were unchanged

Lactation is often associated with reduced anxiety-like behavior, largely mediated by elevated central OXT signaling^93,94^. Consistent with this direction, *LD20* dams showed a trend toward faster re-entry versus *virgin* females, but this did not generalize to other LDB parameters. Crucially, separated mothers (*LD1+19*) did not differ from *LD20* or *virgin* females on any measure, indicating that offspring loss did not produce robust changes in anxiety-related behavior. The absence of locomotor differences across groups further supports that the FST effects were not secondary to altered mobility or general activity. While the lack of pup-derived sensory stimulation in separated mothers could diminish central OXT release -and thereby remove a potential anxiolytic mechanism^95^ -the present data indicates no measurable anxiety-like phenotype in the LDB. Subtle behavioral differences observed in *virgin* females may reflect their period of social isolation, a known driver of heightened anxiety/hyperactivity in rodents.^96^

### Offspring loss increased passive stress-coping and HPA axis reactivity

Our results show that permanent offspring loss exerted a sustained impact on maternal emotional regulation. In the FST, separated mothers consistently exhibited elevated passive stress-coping, as indicated by increased floating time, compared with both *virgin* and *LD20* dams. This outcome replicates and extends previous findings that prolonged maternal separation enhances passive stress-coping in rat dams¹⁰, ¹¹,¹³. The consistent emergence of this phenotype across independent cohorts emphasizes its robustness and supports the validity of this model for probing maternal adaptations to offspring loss. Importantly, locomotor activity did not differ between *LD1+19* and *LD20* dams, confirming that the increase in floating reflects a shift in stress-coping strategy rather than reduced motor capacity or energy availability. In addition, physiological analyses aligned with these behavioral findings. *LD20* mothers displayed the known dampened HPA axis response, consistent with reduced stress reactivity during the postpartum period^97^. By contrast, *LD1+19* mothers exhibited CORT levels similar to *virgin* females, indicating that offspring loss disrupts the typical stress-buffering adaptations of lactation. This reversion to a virgin-like endocrine profile suggests both heightened vulnerability to acute stressors and the absence of maternal neuroendocrine mechanisms that normally confer resilience during caregiving. Whether this shift reflects the acute impact of pup loss on HPA axis regulation or the lack of sustained maternal experience remains an open question. Together, these findings demonstrate that offspring loss induces a dual effect: an increase in passive stress-coping behavior and the reactivation of heightened HPA axis activity. This behavioral–endocrine combination provides a mechanistic framework for understanding how the maternal brain adapts—or fails to adapt—after loss and reinforces the translational relevance of this model.

### Central CRF-R1/2 antagonism, but not OXT-R agonism, reversed passive stress-coping

During lactation, activity of the brain CRF system is normally dampened, providing resilience against stress and allowing mothers to maintain adaptive caregiving responses^98^. However, chronic or severe stress around parturition can disrupt this regulation, leading to heightened CRF signaling and increased vulnerability to stress^99,100^. Furthermore, the CRF system has been implicated in behavioral adaptations following social bond disruption, as shown in prairie voles where partner separation activates CRF-dependent stress responses^101^. In line with this, our results demonstrate that icv administration of the CRF-R1/2 antagonist D-Phe normalized the increased passive stress-coping observed in separated mothers during the FST. These findings indicate that offspring loss induces hyperactivity of the CRF system and that CRF receptor blockade can effectively restore adaptive stress-coping. This interpretation aligns with prior work showing that D-Phe alleviates separation-induced behavioral alterations in both lactating^101^ and male prairie voles¹⁶. Given the known functional interactions between CRF and OXT systems^18,22^, we also investigated whether enhancing OXT signaling could counteract the behavioral phenotype. Previous studies in prairie voles demonstrated that local OXT infusion into the NAcc shell rescues separation-induced behavioral deficits¹⁸, and work in lactating rats highlights OXT’s regulatory role in stress responses. However, in our study central OXT infusion did not modify passive stress-coping in separated mothers. This lack of effect does not exclude the OXT system as a relevant modulatory pathway; rather, it suggests that global central infusion may not capture the region-specific actions of OXT. Indeed, targeted manipulations in regions such as the PVN^102^ or NAcc have produced stronger behavioral effects in other models. Together, our data highlight CRF hyperactivity as a central mechanism in the stress-related consequences of offspring loss, while suggesting that OXT-based interventions may require more precise spatial targeting to be effective.

## Conclusions

This study establishes a translational rat model to investigate the neurobiological consequences of permanent offspring loss. Separated mothers displayed altered stress-coping strategies, elevated HPA axis activity, and structural as well as receptor-level adaptations in limbic brain regions. Specifically, increased OXT-R binding and reduced dendritic spine density were observed in the VMH, identifying this region as a key locus of maternal adaptation to loss. Importantly, pharmacological blockade of CRF-R1/2 signaling restored normal coping behavior, whereas OXT infusion did not, underscoring the CRF system as a primary driver of these outcomes. These findings provide novel insights into the interplay between neuroendocrine signaling, neuroplasticity, and behavioral regulation in the maternal brain. They also suggest that interventions targeting CRF signaling hold particular promise for mitigating the long-term consequences of disrupted maternal experience.

## Acknowledgements

We thank Dr. J. Pawluski and Dr. B. di Benedetto for scientific discussion. We also thank M. Fuchs, A. Havasi, R. Maloumby, M. Schwarz, M. Breitkopf, L. Saller, E. Rocaboy, S. Pena-Pena and T. Gebhardt for excellent technical help.

This study was supported by the Deutsche Forschungsgemeinschaft (DFG), through the Neuroscience Graduate Programme “Neurobiology of Emotion Dysfunctions” (GRK 2174, to O.J.B.) and the grant BO1958/8-2 (to O.J.B.).

## Author contributions

L.D. designed the study, performed the experiments, analyzed data, and wrote the first draft of the manuscript. A.S. designed the study, performed the experiments and revised the manuscript. A.B. performed the experiments. O.J.B. designed the study, secured funding, supervised the project, and revised the manuscript. All authors contributed to and have approved the final manuscript.

## Data availability statement

The data that support the findings of this study are available from the corresponding author upon reasonable request.

